# Ecological Feedback Dimensionality and Trade-Off Shape Govern Cancer Phenotypic Diversification

**DOI:** 10.64898/2026.11.14.766691

**Authors:** Anuraag Bukkuri, György Barabás

## Abstract

Cancer cells face a variety of trade-offs in their microenvironment, from competition vs. colonization to immune evasion vs. rapid proliferation. Whether such trade-offs drive cancer cell populations to a single optimal phenotype or to a stable coexistence of phenotypes depends on the trade-off’s shape and the number of independent ecological feedbacks that cells impose on each other and on their environment. We introduce trade-off and invasion plots (TIPs), a geometric tool from adaptive dynamics that determines the evolutionary outcome from the curvature of a trade-off relative to a pair of invasion boundaries, to cancer and ground our analysis in simple consumer-resource metabolic models. We show that the dimension of the ecological feedback determines whether diversification can occur: a one-dimensional feedback leads to monomorphism, whereas a two-dimensional feedback can promote evolutionary branching to a stable dimorphism. Furthermore, we show that the curvature of the allocation trade-off governs whether a single metabolic generalist (attractor), a stable dimorphism of specialists (evolutionary branching), or a single specialist (repellor) occurs. Both results are cancer-specific instances of long-established principles in theoretical ecology, namely the bound that the dimension of the feedback environment places on coexistence, and the fitness-set geometry relating trade-off curvature to specialization. Our contribution therefore primarily lies in bringing this mature eco-evolutionary machinery to bear on intratumoral heterogeneity.

## 1 Introduction

As the economist Thomas Sowell once proclaimed, “There are no solutions, only trade-offs” [1]. Cancer cells, much like all organisms, cannot optimize across all traits simultaneously: Along the Pareto frontier, a fitness benefit acquired via one trait comes at a fitness cost suffered via another. For instance, a cell exhibiting the Warburg phenotype rapidly converts glucose to ATP, but does so wastefully, trading yield for speed [2, 3, 4]. Analogously, a cell exposed to therapy may enter a dormant state to survive therapy, but does so at the expense of proliferation [5, 6, 7, 8]. Such trade-offs are at the heart of our understanding of cancer as an ecological and evolutionary process [9, 10] and form the basis of innovative evolutionary therapies such as adaptive therapy (which relies on a cost of resistance) [11, 12, 13] and double bind therapies (which rely on resistance to one therapy conferring sensitivity to another) [14, 15, 16].

Trade-offs also contribute to intratumoral heterogeneity [17, 18, 19]. In this paper, we draw on trade-off and invasion plots (TIPs), a tool from geometric adaptive dynamics, to understand the conditions under which trade-offs lead to stable diversification of cancer clones, in contrast to merely selection for an optimal phenotype or divergence to an extreme. The evolutionary impact of a trade-off is governed by the curvature of the trade-off frontier (whether the traits are substitutes or complements) and the dimension of the ecological feedback among cells. Because the fitness of cancer cells depends on their environment, which is shaped by the traits and densities of other cancer cells, evolution is inherently density- and frequency-dependent. Thus, optimization of a fixed fitness metric no longer holds [20, 21].

Given an evolutionarily singular strategy, a trait value for which the selection gradient vanishes, two natural questions arise: 1) Will gradual evolutionary steps move the population towards the singular strategy (convergence stability: CS)? and 2) Is the singular strategy uninvadable by a nearby mutant (evolutionary stability: ES) [21, 22]? This gives rise to four possibilities:

- If a singular strategy has both CS and ES, it is a continuously stable strategy (CSS)–the population converges to a single phenotype, an attractor.
- If a singular strategy is CS but not ES, evolutionary branching will occur–the population initially converges, but then splits, giving rise to a stable dimorphism.
- If a singular strategy is ES but not CS, it is a Garden-of-Eden point–it sits at an uninvadable equilibrium that evolution never reaches. We do not encounter Garden-of-Eden points in the models below, so we will not discuss this further.
- If a singular strategy is neither CS nor ES, it is a repellor–the dynamics move the population away from the singular strategy. Although this may resemble evolutionary branching, it results in a monomorphism rather than a dimorphism.

The box below summarizes key equilibrium concepts from adaptive dynamics that we will refer to throughout the text. For a more comprehensive introduction to adaptive dynamics, we refer the readers to [23, 24]. Which evolutionary endpoint is reached can be ascertained via TIPs [25, 26, 27], an extension of Levins’s geometric comparison of fitness sets and adaptive functions [28] to account for density- and frequency-dependence [29] (Fig. 1). In this paper, we bring TIPs to cancer and foreground trade-offs as a critical factor in promoting intratumoral heterogeneity. To do this, we analyze two related metabolic models to understand when trade-offs lead to a single generalist phenotype (attractor), a stable dimorphism of specialists (evolutionary branching), or a single specialist (repellor). We show that the number of independent ecological feedbacks determines whether stable diversification is possible, and the shape of the trade-off curve determines which regime occurs.

**Figure 1.**
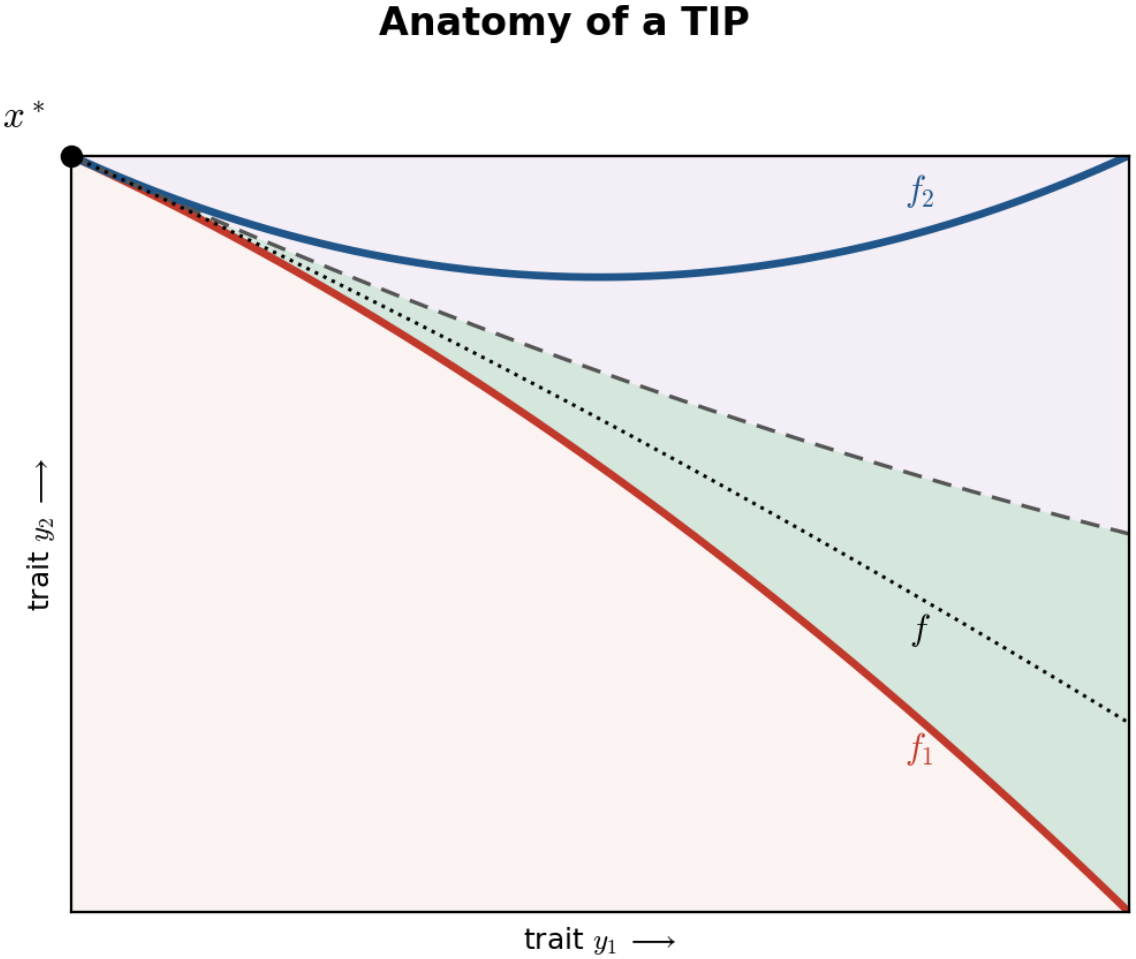
How to read a TIP. The invasion boundaries *f*_1_ (red) and *f*_2_ (blue) are the zero invasion fitness contours, along which a mutant’s fitness equals the resident’s fitness. They meet tangentially at the singular strategy *x*^*\**^, along with the trade-off curve *f* (dotted). *f*_1_ denotes the ES boundary, and the dashed line denotes the mean-curvature CS boundary. The evolutionary outcome is determined by which region the trade-off curve lies in: if it is in the red region, *x*^*\**^ is an attractor and the population converges to it; if it is in the green region, evolutionary branching occurs and the population splits into two coexisting phenotypes; if it is in the purple region, *x*^*\**^ is a repellor and the population evolves away from it, to one extreme of the trade-off (which extreme depends on the initial condition).

Our results below have well-established counterparts in theoretical ecology, and we will highlight the correspondences as we go. For example, the fact that the number of independent regulating factors provides an upper bound to the number of coexisting strategies has first been claimed as a general statement by Levin [30], which has since formed the basis of a rigorous set of theories about not just how many species there could be at most, but how sensitive one should expect any species assemblage to be against external environmental perturbations [31, 32, 33, 34]. Similarly, the idea that a one-dimensional feedback collapses eco-evolutionary dynamics back into ordinary optimization is known variously as the *K*-maximization principle [35], the *R*^*\**^ rule [36], or the pessimization principle [37, 38, 39]. That the curvature of a trade-off decides between generalists and specialists goes back to Levins’s fitness sets [28], subsequently corrected to account for density- and frequency-dependence [29, 40, 25]. Finally, evolutionary branching on two substitutable resources is among the textbook examples of adaptive diversification [41, 42, 43]. Given these well-established results, our aim is not to announce new theory but to make an existing and mature body of eco-evolutionary theory usable by mathematical oncologists, and to show that the questions cancer biologists already ask about intratumoral heterogeneity have precise, previously answered analogues in community ecology.

## 2 A Primer on TIPs

**Key Equilibrium Concepts from Adaptive Dynamics**

**Invasion fitness** *s*_*x*_(*y*). Long-term per-capita growth rate of a rare mutant clone *y* in the environment set by a resident clone *x*.

**Singular strategy** *x*^*\**^. A trait value at which the selection gradient vanishes and directional selection stops.

**Convergence stability**. Successive small mutations drive the population towards *x*^*\**^.

**Evolutionary stability**. *x*^*\**^ is a fitness peak; no nearby mutant can invade.

**Branching point**. A singular strategy that is convergence stable but not evolutionarily stable: the population reaches it, then diversifies into two phenotypes that coexist (stable heterogeneity).

In our models, a clone is described by two traits, *y*_1_ and *y*_2_, which capture resource allocation between rate and yield of energy metabolism or between transporters for different nutrients. A trade-off curve is imposed such that fixing one trait determines the other, i.e., *y*_2_ = *f* (*y*_1_). This is analogous to defining the Pareto frontier in economic models, intentionally eliminating dominated options to simplify analysis [44]. In this way, each clone fundamentally depends on a single trait *y*. Classical invasion analysis from adaptive dynamics asks: If we fix one clone as resident at its corresponding ecological equilibrium, can a rare nearby mutant invade the population? The answer depends on the invasion fitness *s*_*x*_(*y*), which captures the long-term per capita growth rate of the rare mutant *y* in an environment set by the resident *x*. If *s*_*x*_(*y*) *>* 0, the mutant invades the population; if *s*_*x*_(*y*) *<* 0, the mutant goes extinct. An invasion boundary for *x* can be found by sweeping across mutant trait values and tracing the curve for which *s*_*x*_(*y*) = 0.

A reader familiar with adaptive dynamics will notice the similarities between TIPs and pairwise invasibility plots (PIPs), the standard tool to visualize invasion fitness [21]. These two techniques capture the same invasion fitness, *s*_*x*_(*y*), differing only in the axes upon which they choose to graph it. Namely, PIPs plot invasion fitness in resident-mutant space: the diagonal *y* = *x* captures a zero invasion fitness boundary (*s*_*x*_(*x*) = 0) and a second zero invasion fitness contour captures the singular strategies. Depending on the sign patterns around the singular strategy, properties of convergence and evolutionary stability can be ascertained. Of relevance here, branching occurs in regions of mutual invasibility when *s*_*x*_(*y*) *>* 0 and *s*_*y*_(*x*) *>* 0. TIPs on the other hand plot the invasion boundaries and trade-off frontier in trait space–mutual invasibility and evolutionary branching are possible when the invasion boundaries separate. For questions in which trade-offs are central, TIPs provide a distinct advantage over PIPs. Specifically, the latter implicitly handles trade-offs within the fitness functions; thus, exploring various aspects of the trade-off function would require one to generate separate PIPs for each case. However, TIPs foreground trade-offs by capturing them as independent curves alongside fixed invasion boundaries. This way, exploring trade-off function properties and how they impact evolutionary outcomes can be done with ease.

As described in [25], a TIP analyzes the geometry of the trade-off at the singular strategy through three curves: the invasion boundary *f*_1_ for the resident *x*, the invasion boundary *f*_2_ for the mutant *y*, and the trade-off curve *f*, where *y*_2_ = *f* (*y*_1_). Since *s*_*x*_(*x*) = 0 by definition, all three curves are tangent at the singular strategy *x*^*\**^ (Fig. 1). At this point, the selection gradient vanishes, and directional selection stops.

The challenge then is to determine what happens at this singularity: Does evolution converge to *x*^*\**^, repel from it, or branch into two coexisting phenotypes? The answer depends on the relation of the three curves outlined above. As shown in [25], when fitness increases in the second trait at the singularity (*∂s/∂y*_2_ *>* 0), which holds in both our models, the outcome is classified as follows:

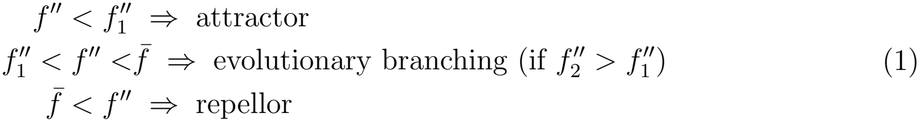

where 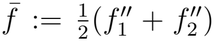 is the mean curvature. Note that branching can occur only if there is a separation of the two invasion boundaries, i.e., 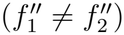. As derived in [25], this separation is governed by the identity

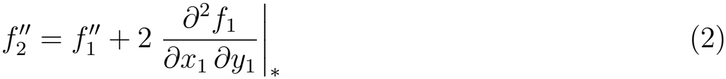

Thus, branching requires that the mixed partial 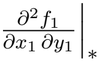 is nonzero. These relations among curvatures can be evaluated either analytically or numerically via TIP plots (Fig. 1). In the following examples we provide both, as each offers its own advantage: analytical calculation determines the outcome as exact conditions on the model parameters, whereas geometric visualization shows all three regimes together and where a given trade-off sits among them at a glance, making the structure behind those conditions visible.

## 3 Results

To investigate how TIPs may be used to understand intratumoral heterogeneity, we focus on two related metabolic models. In both, a cancer clone’s traits determine how it allocates a fixed energetic budget between two competing metabolic functions, *u*_1_ and *u*_2_, which represent either metabolic rate vs. yield (Example 1) or investment into transporters for distinct nutrients (Example 2). The set of non-dominated trait pairs lies on the Pareto frontier

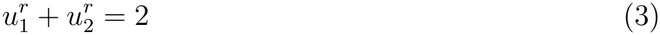

The parameter *r* captures the curvature of the frontier, in which higher values of *r* correspond to more substitutable traits. With this setup, we are now ready to construct and analyze our specific metabolic models. For each example, we will provide both TIPs and corresponding eco-evolutionary simulations capturing the reachable regimes. The simulations we perform are mutation-limited, trait substitution sequences, adaptive dynamics in style. For simplicity, we track a scalar allocation parameter *v* := *u*_1_*/*(*u*_1_ + *u*_2_) *∈* (0, 1)–a balanced generalist will thus have *v* = 0.5, with *u*_1_ and *u*_2_ specialists at *v* = 1 and *v* = 0, respectively. At each time step, a rare mutant with a trait value randomly sampled from a narrow Gaussian distribution about the resident trait is introduced into the population. Its invasion fitness is calculated in the ecological equilibrium set by the resident. If its invasion fitness is negative, it is removed from the population. Assuming invasion implies substitution, if the mutant has a positive invasion fitness, it replaces the resident in the population and a new ecological equilibrium is computed. However, if the resident and the mutant are mutually invasible, i.e., *s*_*x*_(*y*) *>* 0 and *s*_*y*_(*x*) *>* 0, they coexist as a protected dimorphism and evolutionary branching occurs. By changing *r*, we can change the location of the trade-off curve and move through the attractor, branching, and repellor regions of the TIP.

### 3.1 Example 1: Rate vs. Yield of Energy Metabolism

One metabolic trade-off we can consider is that between the rate and the yield of energy metabolism. For instance, a cell adopting a Warburg phenotype may run glycolysis, which rapidly converts glucose to ATP but is wasteful. Alternatively, a cell may run oxidative phosphorylation, a metabolic program which is much more efficient, but converts glucose to ATP more slowly [2, 3, 4]. The question then arises: Does a population with a shared glucose resource adopt a singular metabolic phenotype or split into two populations, with fast and efficient specialists?

To address this, we develop a basic consumer-resource model. We let *N* be the density of a cancer clone and *R* be the shared glucose concentration. Each cell takes up glucose at rate *JR*, converts it to biomass with yield *e*, and pays a constant cost of maintenance *m*. Glucose is supplied at a constant rate *I*, decays at a per-unit rate *w*, and is consumed by the cancer population at rate *JRN*. We formalize this as:

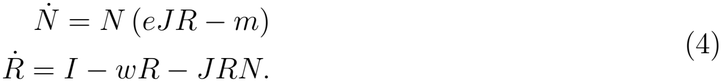

Traits *e* and *J* satisfy the Pareto frontier (3): *e*^*r*^ + *J*^*r*^ = 2, allocating a fixed energetic budget between rate and yield. By symmetry, the singular strategy is *e*^*\**^ = *J* ^*\**^ = 1; the frontier has slope *f* ^*′*^(*J* ^*\**^) = *−*(*e*^*\**^*/J* ^*\**^)^1*−r*^ = *−*1 and curvature *f* ^*′′*^(*J* ^*\**^) = 2(1 *− r*). At the non-trivial equilibrium, a resident with trait pair (*e*_*x*_, *J*_*x*_) has a per-capita birth rate that equals its maintenance cost such that

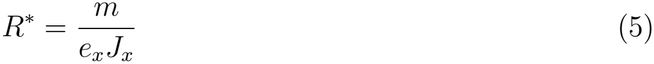

This captures the ecological feedback that the resident exerts on competing clones. A rare mutant (*e*_*y*_, *J*_*y*_) that is introduced into the environment set by the resident grows at rate

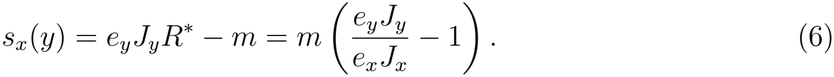

#### Proposition 1

*At the singular strategy, the two invasion boundaries coincide, so branching cannot occur. x*^*\**^ *is an attractor if r >* 0 *and is a repellor if r <* 0.

*Proof*. Setting (6) to zero gives the invasion boundary

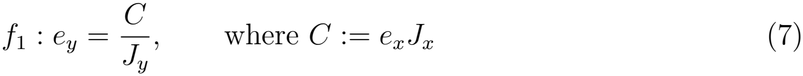

The resident thus impacts the boundary only through the single scalar *C*, a one-dimensional ecological feedback. Along the trade-off, *C* = *f* (*J*_*x*_)*J*_*x*_, so *C*^*′*^(*J* ^*\**^) = *f* ^*′*^(*J* ^*\**^)*J* ^*\**^ + *f* (*J* ^*\**^) = 0. As *f*_1_ depends on the resident only through *C, ∂*^2^*f*_1_*/∂J*_*x*_ *∂J*_*y*_ = *−C*^*′*^(*J* ^*\**^)*/J* ^*\**2^ = 0. By Equation 2, 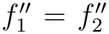–the invasion boundaries are superimposed^*\**^ and branching cannot occur. To determine when the singular strategy is an attractor or a repellor, we must compare *f* ^*′′*^(*J* ^*\**^) = 2(1*−r*) and 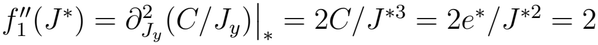. Thus, by classification 1, *x* is an attractor when *r >* 0 and is a repellor when *r <* 0.

Proposition 1 shows us that a monomorphic outcome is guaranteed. We visualize the TIP plot and the corresponding eco-evolutionary simulations in Fig. 2 below. As we can see, the invasion boundaries *f*_1_ and *f*_2_ are superimposed, leaving no room for evolutionary branching. The attractor phenotype is generated by setting *r* = 0.5 and occurs when 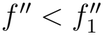–all trajectories converge to the generalist phenotype at *v* = 0.5. The repellor phenotype is generated by setting *r* = *−*1 and occurs when 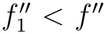–clones that start below *v* = 0.5 become yield specialists, whereas clones whose trajectories start above *v* = 0.5 become rate specialists.

**Figure 2.**
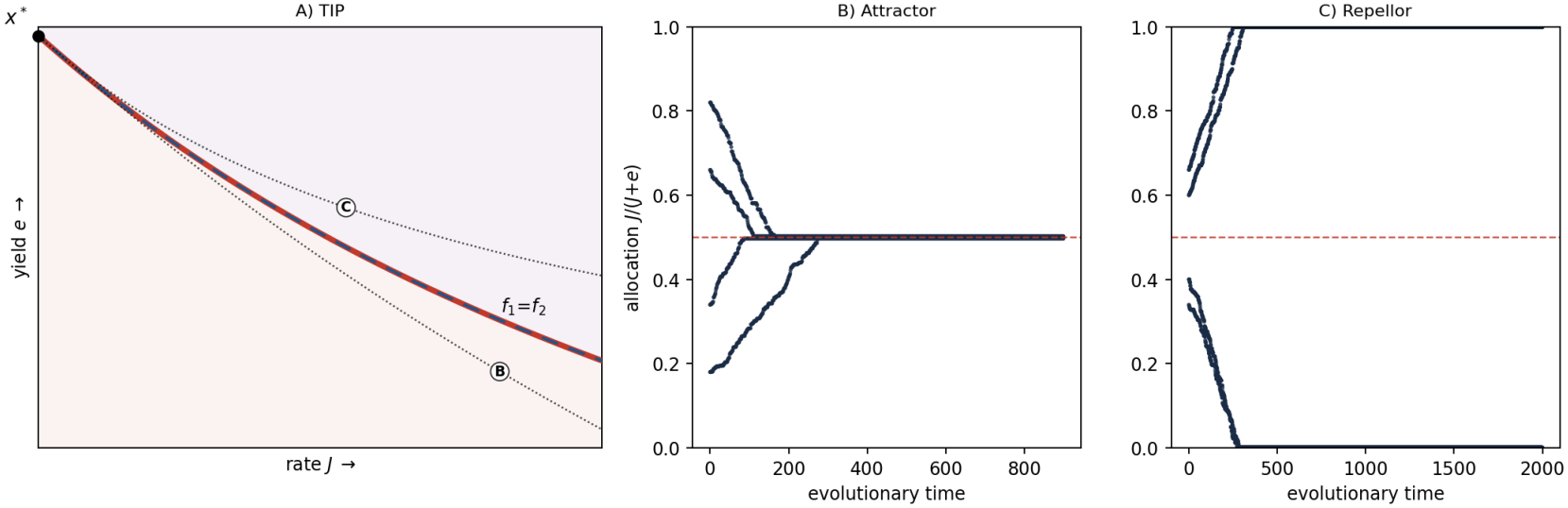
Rate–yield metabolism. (A) TIP: a one-dimensional ecological feedback leaves the invasion boundaries superimposed, with no room for evolutionary branching. An attractor regime exists below *f*_1_ and a repellor regime exists above it. Dotted curves denote the trade-off used in panels B & C. (B,C) Trait-substitution simulations: all trajectories converge to *v* = 0.5 for the attractor (B). For the repellor (C), trajectories that start above (below) *v* = 0.5 diverge to *v* = 1 (*v* = 0).

This outcome is not an artifact of our particular parameterization. It is the cancer-specific instance of one of the oldest results in resource competition theory. With a single limiting resource, the resident’s entire influence on every other clone is transmitted through the scalar *R*^*\**^, and invasion fitness (6) is a strictly increasing function of that one number. Selection therefore acts exactly as if it were maximizing *eJ* (or equivalently, minimizing *R*^*\**^). The clone able to persist at the lowest glucose concentration excludes all others. This is Tilman’s *R*^*\**^ rule [36], together with its evolutionary counterpart, the pessimization principle, which holds that when the feedback environment is one-dimensional, density- and frequency-dependence cannot produce anything that a plain optimization argument would miss [37, 38, 39]. One resource means one winner, regardless of the shape of the tradeoff. The curvature can only decide which single phenotype wins. The vanishing mixed partial in the proof above is the geometric signature of this fact: with a single regulating factor the resident cannot tilt the invasion boundary, so the two boundaries have nowhere to separate [45].

### 3.2 Example 2: Transporter Investment for Distinct Nutrients

Another metabolic trade-off we can consider is investment in the uptake of distinct nutrients. Cancer cells, much like organisms in the natural world, use a variety of nutrients to fuel their growth and must invest in the production and maintenance of transporters on their cell surface to take up these nutrients. In this section, we consider what happens when cancer cells are given two nutrients and must allocate a fixed energy budget to obtain each resource. We let *u*_1_ and *u*_2_ represent investment into uptake of resources 1 and 2, respectively. For simplicity, we assume that uptake capacity is a linear function of trait value, although the results hold for diminishing returns. The model structure is analogous to the one presented in the previous section:

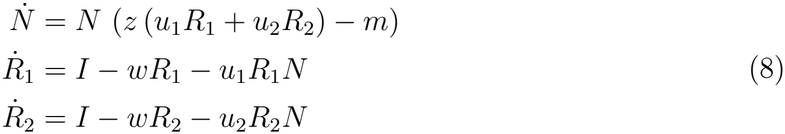

The investments satisfy the same Pareto frontier (3): 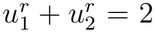, so by symmetry the singular strategy is again 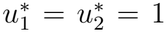. At the resident equilibrium, a mutant with traits (*u*_1,*y*_,*u*_2,*y*_) has the invasion fitness

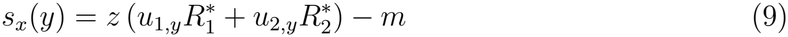

#### Proposition 2

*At the singular strategy, the two invasion boundaries separate, so branching is possible: x*^*\**^ *is an attractor when r >* 1, *a branching point when wm/*(2*Iz*) *< r <* 1, *and a repellor when r < wm/*(2*Iz*).

*Proof*. Setting (9) to zero gives the invasion boundary

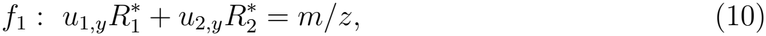

a straight line in the mutant’s traits with slope 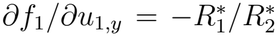, so 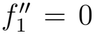. At the singular point, the two resources coincide so 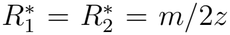 (for simplicity, we will just call this *R*^*\**^). Implicit differentiation of the resident equilibrium (8) along the trade-off then gives 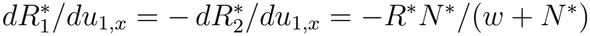, so

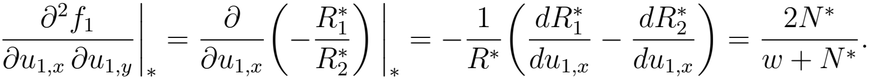

By (2), 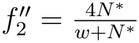 and since 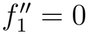, the boundaries separate and a regime for evolutionary branching opens between the ES boundary 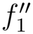 and the mean 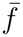. Since *f* ^*′′*^ = 2(1 *− r*), by classification (1), the attractor/branching edge 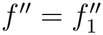 gives *r* = 1, and the branching/repellor edge 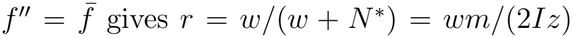. Thus *x*^*\**^ is an attractor when *r >* 1, a branching point when *wm/*(2*Iz*) *< r <* 1, and a repellor when *r < wm/*(2*Iz*).

Proposition 2 shows that the addition of a second independent resource allows for a two-dimensional ecological feedback, which provides the conditions for evolutionary branching. It is worth noting that the branching window widens as resource influx or yield increases and narrows as maintenance cost or natural resource degradation rises. In other words, more resource-rich or efficient environments promote conditions for tumor diversification. We visualize the TIP plot and the corresponding eco-evolutionary simulations in Fig. 3. As we can see, there is a clear separation between the invasion boundaries *f*_1_ and *f*_2_, leaving room for evolutionary branching. The attractor phenotype is generated by setting *r* = 1.5 and occurs when 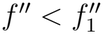–all trajectories converge to the generalist phenotype at *v* = 0.5. Evolutionary branching is generated by setting *r* = 0.5 and occurs when 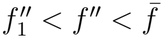–trajectories approach *v* = 0.5, before branching into nutrient 1 (*v* = 1) and nutrient 2 (*v* = 0) specialists. The repellor phenotype is generated by setting *r* = *−*1 and occurs when 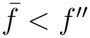–clones that start below *v* = 0.5 become nutrient 2 specialists, whereas clones whose trajectories start above *v* = 0.5 become nutrient 1 specialists.

**Figure 3.**
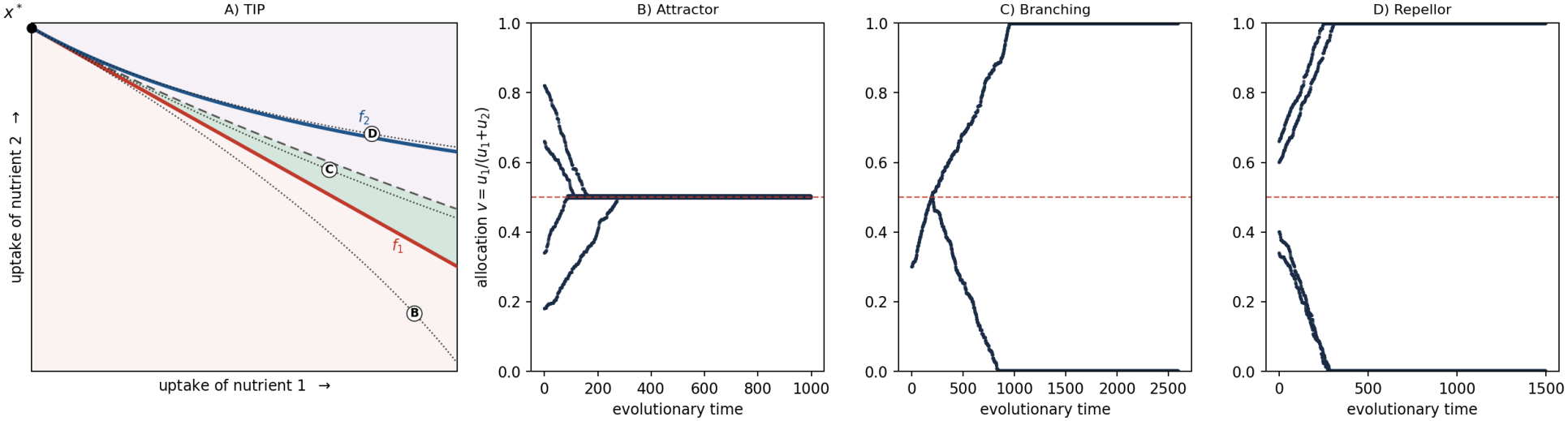
Investment in uptake of two nutrients. (A) TIP: a two-dimensional ecological feedback separates the invasion boundaries, leaving room for evolutionary branching. All three regimes (attractor, branching, repellor) now exist. Dotted curves denote the trade-off used in panels B-D. (B-D) Trait-substitution simulations: all trajectories converge to *v* = 0.5 for the attractor (B). Under evolutionary branching (C), trait trajectories approach the generalist strategy before diversifying into nutrient 1 and 2 specialists. For the repellor (D), trajectories that start above (below) *v* = 0.5 diverge to *v* = 1 (*v* = 0).

Here again, ecological theory arrived first. Structurally, Example 2 is the same as MacArthur’s consumer-resource model with two resources [31, 36], and the result that a consumer evolving on two substitutable resources can branch into two specialists is one of the standard illustrations of evolutionary branching [41, 42, 43]. The sequence of outcomes we recover (a generalist when the resources are highly substitutable, branching at intermediate substitutability, and divergence to a single specialist when they are complements) reproduces the classical weak versus strong trade-off dichotomy of Levins’s fitness sets [28], supplemented by the intermediate branching window that only appears once density- and frequency-dependence are accounted for [29, 46, 47, 48]. Egas and colleagues made this point for herbivorous arthropods: it is the structure of the trade-off, not the mere availability of two resources, that determines whether specialists coexist [49]. A TIP renders that same distinction visually, as the position of the trade-off curve within the wedge opened between *f*_1_ and 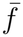.

## 4 Conclusion

The importance of intratumoral heterogeneity for a variety of oncological phenomena, from cancer initiation and diversification to therapeutic response and metastatic progression, cannot be overstated. Understanding why stable diversification occurs in tumors is the first step to designing ecologically and evolutionarily enlightened therapies for durable responses. In this paper, we introduced the idea of trade-off and invasion plots (TIPs), a geometrical approach to adaptive dynamics that foregrounds the role of trade-offs, to the cancer community. We explained how to construct and interpret TIPs, both as plots and through the curvature conditions that underlie them, to assess whether trade-offs lead to stable diversification. We then showed that the dimension of the ecological feedback determines the plausibility for evolutionary branching to occur, and that the shape of the trade-off curve determines whether the singular strategy manifests as an attractor, a repellor, or leads to evolutionary branching. Although we grounded our framework in a simple class of consumer-resource metabolic models, our results apply more broadly across cancer trade-offs: competition-colonization, go-or-grow, immune evasion-proliferation, and therapy resistance-cycling can all be analyzed using the TIP framework we presented.

This provides a testable prediction: a cancer cell population limited by a single resource should evolve towards one optimal metabolic strategy, whereas a population with two independently limiting resources may stably diversify into coexisting specialists, assuming that the two uptake systems are imperfect substitutes. The strength of substitutability governs whether this diversification occurs in practice. Therapeutically, if heterogeneity is maintained by a high-dimensional ecological feedback, interventions that collapse a multi-resource environment into one (e.g., by starving a nutrient or preventing a cross-feeding exchange), could promote homogenization and improve the efficacy of a subsequent therapy.

Framed this way, what is new here is the target rather than the content. The bounds on coexistence or the curvature conditions for evolutionary branching are parts of theoretical ecology, and have been for some time [30, 20, 21, 50, 32]. The important recognition is that intratumoral heterogeneity poses the same questions community ecologists have long asked about coexistence [51], and therefore inherits both the same answers and the same constraints: heterogeneity not underwritten by a sufficient number of independent regulating factors is not stable, and no amount of mutational supply will make it so. Cancer ecoevolutionary theory need not rediscover these results one model at a time. Once tumors are treated as ecological communities in earnest [9, 52, 53], the mature machinery of coexistence theory becomes available to oncology, which TIPs are one part of.

Although TIPs were introduced decades ago in evolutionary ecology, they remain entirely unused in cancer evolutionary ecology. In light of the growing number of eco-evolutionary mathematical models of cancer and innovative therapeutic strategies that rely on trade-offs among cancer cell clones, we hope that this paper provides a valuable technique in the toolbox of mathematical oncologists interested in cancer evolution.

